# Rapid engineering of SARS-CoV-2 therapeutic antibodies to increase breadth of neutralization including XBB.1.5 and BQ.1.1

**DOI:** 10.1101/2023.01.25.525589

**Authors:** Kevin C. Entzminger, Jonathan K. Fleming, Paul D. Entzminger, Lisa Yuko Espinosa, Alex Samadi, Yuko Hiramoto, CJ Okumura, Toshiaki Maruyama

**Author notes:** Correspondence should be addressed to T.M.

## Abstract

An antibody panel that broadly neutralizes currently circulating Omicron variants was obtained by *in vitro* affinity maturation using phage display. Starting from a single parent clone, antibody engineering was performed in iterative stages in real time as variants emerged using a proprietary technology called STage-Enhanced Maturation (STEM). Humanized from a rabbit antibody, the parent clone showed undetectable neutralization of later Omicron variants, while an early stage IgG possessing only an engineered light chain potently neutralizes some BA.2 but not BA.4/BA.5 lineage variants. However, the final heavy and light chain engineered mAbs show potent neutralization of XBB.1.5 and BQ.1.1 by surrogate virus neutralization test, and biolayer interferometry shows pM K_D_ affinity for both variants. Our work not only details novel therapeutic candidates but also validates a unique general strategy to create broadly neutralizing mAbs to current and future SARS-CoV-2 variants.

---

During the three years since the outbreak of COVID-19, the SARS-CoV-2 virus has proven exceptionally adept at mutating to evade the immune response^1^. The emergence of the first Omicron variant BA.1, containing many more mutations compared to earlier variants, dramatically increased the susceptibility of even previously vaccinated or infected individuals^2^. Later Omicron sublineages BA.2 (from which XBB.1.5 is derived) and BA.4/5 (from which BQ.1.1 is derived) have continued to evolve additional immune-evading mutations, further increasing the likelihood of breakthrough infections^1,3^. At the time of writing, January 2023, XBB.1.5 and BQ.1.1 are dominant circulating strains, comprising 76% of all infections (cdc.gov). As a master of immune evasion, XBB.1.5 has been deemed the most transmissible variant yet^4^ and has the potential to dominate other variants worldwide.

Antibody therapeutics are the standard of care for at-risk populations who are immunocompromised and thus susceptible to adverse COVID infection. Yet antibody therapeutics approved for treatment of COVID-19 have lost efficacy soon after new strains emerged. BA.1 significantly reduced the potency of first-generation antibody therapeutics such as REGEN-COV (Casirivimab-imdevimab)^1^. BQ.1.1 resists neutralization by single or cocktail mAb therapies including Sotrovimab, Bebtelovimab, Bamlanivimab-etesevimab, and Evusheld (Cilgavimab-tixagevimab)^5,6^. XBB.1.5 has been shown to similarly evade neutralizing antibodies^3^. There has been a call for novel, broadly active mAbs urgently needed for prophylactic and/or therapeutic treatment in patients at high risk^5,7,8^, especially given the danger associated with reinfection^9^. No currently US FDA approved antibody therapeutics potently neutralize the latest Omicron variants BQ.1.1 and XBB.1.5.

Existing strategies to engineer neutralizing therapeutic mAbs typically rely on antibody isolation from infected or vaccinated individuals or from immunized humanized mice^10–12^. However, this process is poorly adapted for rapidly evolving targets such as SARS-CoV-2, where the lead discovery phase must be repeated each time a new variant emerges. Instead, we devised an approach that evolves antibody neutralization breadth in real time as the virus itself evolves. Starting from a humanized lead candidate that showed strong neutralization of the Wuhan-Hu-1 strain, we created six separate CDR-targeted libraries and selected the libraries on Wuhan-Hu-1 spike trimer to create diverse CDR pools. These CDRs were then randomly paired and iteratively selected on SARS-CoV-2 variants as they emerged, resulting in a final panel of seven clones that strongly neutralize all Omicron variants including XBB.1.5 and BQ.1.1. This study not only details potential high-value therapeutic mAbs but also a general strategy to elicit broadly neutralizing mAbs for SARS-CoV-2 or other viruses.

## Results

### Therapeutic mAbs were engineered in real time as SARS-CoV-2 variants emerged

Soon after the onset of the COVID-19 pandemic, we used our rabbit discovery platform, which pairs rabbit immunization with Fab-phage display, to create a therapeutic lead candidate mAb. Rabbits were immunized with Wuhan-Hu-1 receptor-binding domain (RBD), and phage libraries were selected on Wuhan-Hu-1 spike protein trimer. Fab neutralization assay was used to identify the potent clone C-A11. We then grafted CDRs onto a human framework to create the humanized clone hN2Y, which strongly neutralized the Wuhan-Hu-1 strain. As evasive SARS-CoV-2 variants emerged, many groups attempted to identify novel antibodies from vaccinated or infected patient samples^13,14^. We instead employed our STage-Enhanced Maturation (STEM) platform to broaden neutralization potency of hN2Y to SARS-CoV-2 variants as they emerged in real time (**Fig. 1**).

**Fig. 1.**
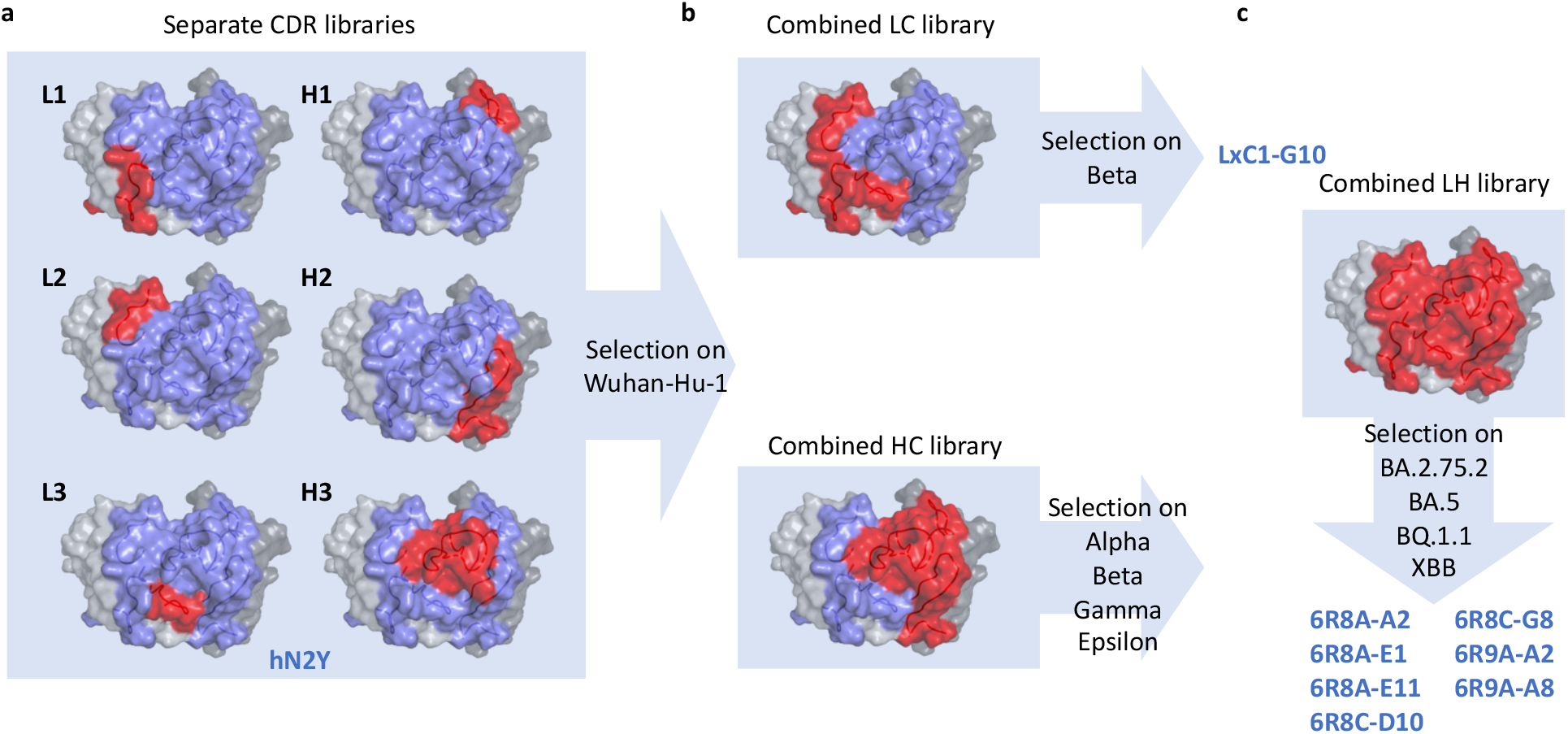
Antibody engineering strategy. **a**, Rabbit immunization with Wuhan-Hu-1 SARS-CoV-2 RBD was paired with phage display to identify a potent neutralizing clone; this mAb was humanized to create the lead candidate hN2Y. Single CDR libraries were constructed and selected on Wuhan-Hu-1 spike trimer to create a pool of functional CDR sequences. **b**, Pre-selected CDR pools were amplified from phage and combined by overlap PCR to create separate combined light and combined heavy chain libraries. The light chain library was further selected on the Beta variant, resulting in the engineered clone LxC1-G10, which showed potent neutralization of BA.1 and was thoroughly characterized by the Coronavirus Immunotherapy Consortium (CoVIC) at La Jolla Institute of Immunology^29^. The heavy chain library was selected on a combination of Alpha, Beta, Gamma, and Epsilon variants to broaden neutralization activity. **c**, Pre-selected light and heavy chain libraries were then paired to create a single combined library. This library was selected on late-stage Omicron variants including BA.2.75.2, BA.5, BQ.1.1, and XBB. Top clones resulting from this library are characterized in detail below and show broad, potent neutralization of all SARS-CoV-2 variants to date.

In traditional humanization, animal-derived CDRs are grafted into human framework germline genes. Then, further engineering is performed using various methods to obtain the best lead candidate for therapeutic mAb development^15–21^. These methods suffer from notable setbacks including (1) the inability to easily sample mutations that work cooperatively within or across CDRs such as in saturation mutagenesis and CDR walking, (2) introduction of unwanted framework mutations such as in error prone PCR, (3) lack of secondary selection pressure when combining beneficial mutations such as in parallel CDR optimization^15^, and (4) almost all omit selection pressure for well-behaved clones. In contrast, our *in vitro* STEM platform is designed to sample the broadest sequence diversity across all CDRs including mutations that work cooperatively and incorporates developability filters during selection.

Single CDR mutant libraries of hN2Y were first created (**Fig. 1a**). CDR libraries were designed based on observed amino acid frequencies for characterized human antibodies possessing a similar germline and predicted CDR canonical structure usage to both further humanize CDRs and to avoid introduction of sequence liabilities. By using phage display, each CDR library can be sampled at sizes up to 2E+10. These separate CDR libraries were first selected on Wuhan-Hu-1 spike trimer to eliminate non-binders and to isolate a large pool of diverse clones. Then, CDR pools from the selected phage populations were amplified and randomly paired by overlap PCR to create secondary combined light chain and combined heavy chain libraries (**Fig. 1b**). To increase neutralization breadth, the combined light chain library was selected on the Beta variant, and the combined heavy chain library was selected separately on Alpha, Beta, Gamma, and Epsilon variants. We identified an exceptional candidate from the combined light chain library, LxC1-G10, which showed good neutralization of BA.1 and BA.2. After the emergence of later BA.2 lineage Omicron variants, we again used our platform to further improve potency. Light and heavy chains were separately amplified from the pre-selected phage pools and further combined to create a single library (**Fig. 1c**). The library was selected on BA.5, BA.2.75.2, BQ.1.1, and XBB variants, all of which possess mutation at F486, which we suspected as a major culprit causing LxC1-G10’s loss of activity. A diverse set of seven highly potent, broadly neutralizing IgGs (6R8/6R9 clones) was identified following 8 or 9 rounds of panning and is characterized below.

### *In vitro* neutralization assay shows broadly neutralizing mAbs

Neutralization titers for sera or monoclonal antibodies measured using surrogate virus neutralization test (sVNT), which assays blocking of recombinant spike trimer to recombinant ACE2 protein by ELISA, shows strong correlation with results from pseudovirus or live virus neutralization tests^22^. We thus used high-throughput sVNT to quantify neutralization potency of mAbs across SARS-CoV-2 variants. In this assay, recombinant human Fcγ-tagged ACE2 receptor is immobilized while IgGs are pre-incubated with spike trimer before addition to the blocked ACE2 coated wells. ACE2-bound spike trimer is detected by affinity tag using ELISA, and IC_50_ values are derived. IgGs tested include the original neutralizing rabbit mAb C-A11, the parent humanized clone hN2Y, LxC1-G10 possessing mutations across light chain CDRs, and 6R8/6R9 clones possessing mutations across both heavy and light chain CDRs.

Notably, humanization of the rabbit antibody did not significantly reduce efficacy, as hN2Y retained similar neutralization potency for Wuhan-Hu-1 and Delta variants compared to C-A11 (**Fig. 2** and **Table 1**). However, both IgGs showed significantly reduced neutralization of the original Omicron variant BA.1. LxC1-G10, engineered as described above, contains mutations within CDRs L1, L2, and L3 compared to hN2Y. Neutralization of early Omicron variants by LxC1-G10 was significantly improved to low ng/mL IC_50_ values for BA.1, BA.2, BA2.75, BA.2.3.20, and BN.1. However, all three IgGs lost efficacy against variant BA.2.75.2, which is notably different compared to these others due to the R346T and F486S mutations. BA.5 and BQ.1 lack the R346T mutation but possess a F486V mutation, and LxC1-G10 does not significantly neutralize these variants, suggesting that mutation at F486 is the most significant contributor affecting loss of LxC1-G10 activity.

**Fig. 2.**
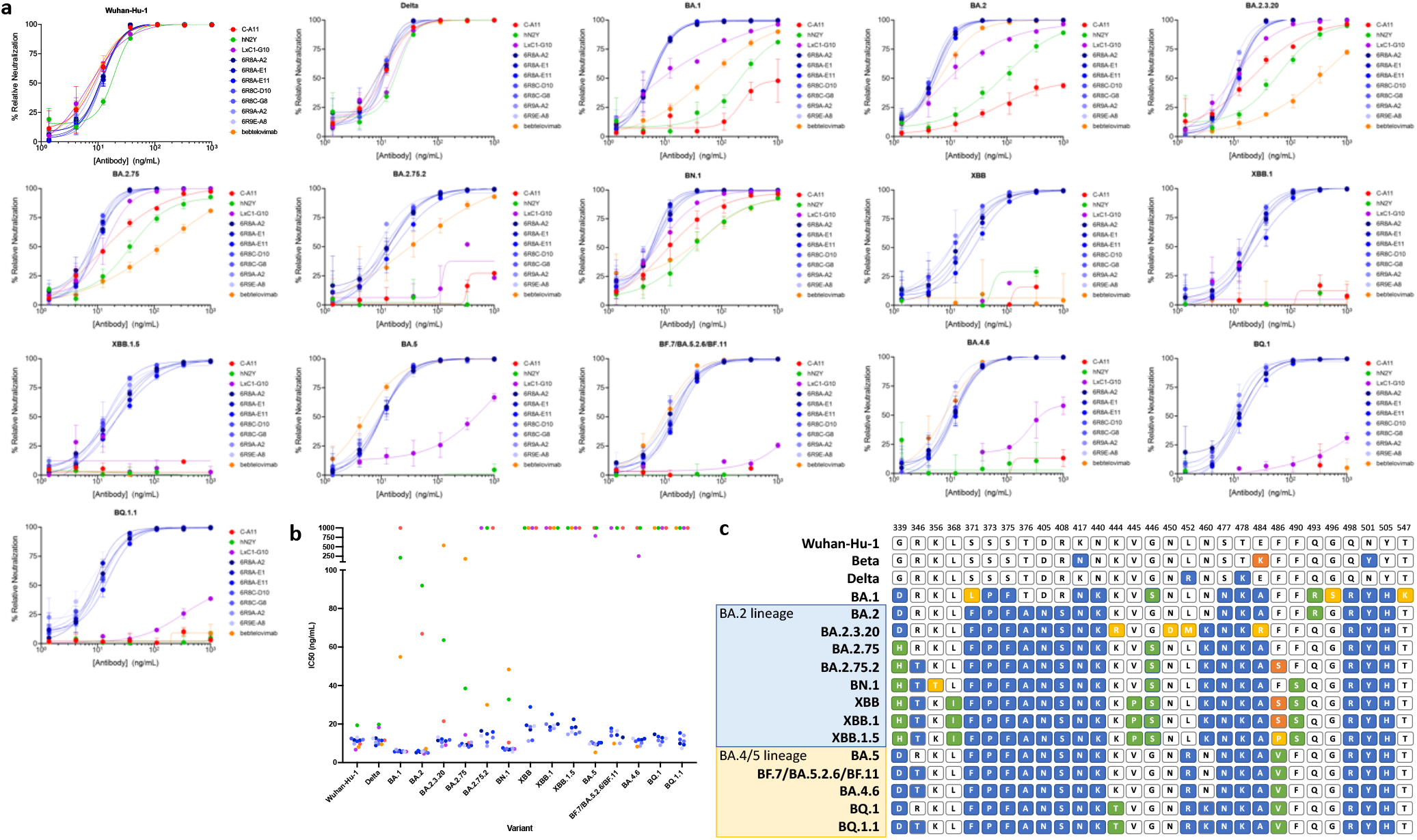
Broad SARS-CoV-2 neutralization potency for engineered IgGs. **a**, Purified IgGs were tested by surrogate virus neutralization test (sVNT)^22^. Wells were coated with recombinant ACE2-Fc, and His-tagged spike trimer protein was mixed with serially diluted mAb prior to addition. Bound spike trimer was detected by anti-His tag antibody. All experiments were performed in triplicates. **b**, Inhibition of binding (IC_50_ ng/mL) was calculated by curve fitting. Non-neutralizing clones are plotted at 1000 ng/mL IC_50_. 6R8/6R9 clones (blue) showed highly potent and broad neutralization of all Omicron variants tested including XBB.1.5 and BQ.1.1. LxC1-G10 (purple) light chain engineered clone showed improved binding compared to lead humanized candidate hN2Y (green) or parental rabbit clone C-A11 (red), though it lost potency for later Omicron variants. Bebtelovimab (orange) showed no neutralization of XBB, XBB.1, or BQ.1.1. **c**, Table of RBD residues modified in the different SARS-CoV-2 variants tested compared to the Wuhan-Hu-1 strain. Residues are colored by mutation.

**Table 1.**
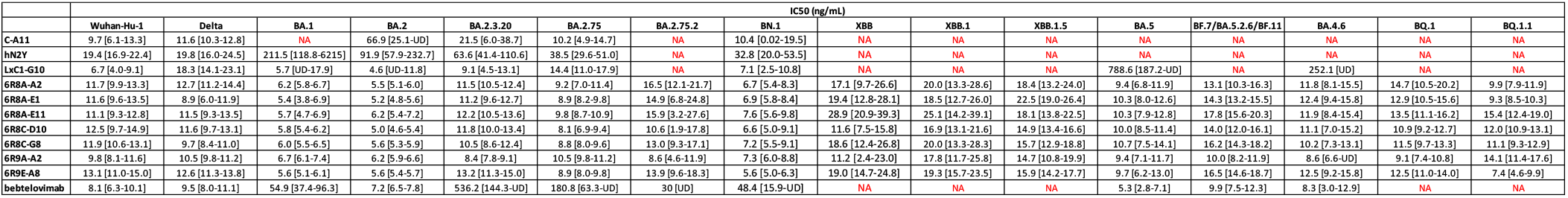
IC_50_ values derived from sVNT data. Calculated IC_50_ values from sVNT assay are provided including 95% confidence intervals. 6R8/6R9 clones show low ng/mL IC_50_ values for all SARS-CoV-2 variants tested. NA, no significant neutralization activity (>1000 ng/mL); UD, undefined.

To recover efficacy, as described above, we engineered a panel of broadly neutralizing antibodies that show strong neutralization activity against all Omicron variants to date including BA.1, BA.2, BA.2.75, BA.2.75.2, BA.2.3.20, BA.4.6, BA.5, BN.1, BQ.1, BQ.1.1, BF.7 (BA.5.2.6/BF.11), XBB, XBB.1, and XBB.1.5. To our knowledge, these are the most broadly neutralizing antibodies characterized to date against the latest Omicron variants of SARS-CoV-2. Compared to both hN2Y and LxC1-G10, these IgG clones possess mutations across CDRs L1, L2, L3, H1, and H3.

Bebtelovimab is a therapeutic SARS-CoV-2 antibody developed by Eli Lilly and first given Emergency Use Authorization by the US FDA in February 2022, though the agency revoked authorization in late November 2022 due to an inability to neutralize variants BQ.1 and BQ.1.1. In our sVNT assay, Bebtelovimab showed no efficacy against these variants or against XBB, XBB.1, or XBB.1.5, matching expected results. S728-1157 clone was developed as a broadly neutralizing antibody against Delta and BA.1^23^. In our sVNT, S728-1157 additionally neutralized BN.1, BA.2, and BA5 but failed to neutralize BA.2.75.2, XBB, XBB.1.5, BQ.1, or BQ.1.1 (**Fig. S1**). Another previously-characterized broadly neutralizing antibody S2X324, identified from human plasma^14,24^, showed strong neutralization of BA.2, BA.2.75.2, and BA.5 in our sVNT but again failed to neutralize XBB.1.5 or BQ.1.1 (**Fig. S2**). In contrast, our engineered IgGs neutralized all SARS-CoV-2 variants tested and all currently circulating variants to date.

### Binding affinity correlates with neutralization potency

Biolayer interferometry (BLI) was performed by capturing biotinylated spike trimer to a streptavidin-coated sensor surface to monitor binding of serially diluted IgGs, with traces globally fit to 1:1 binding models to derive kinetic and affinity values. The humanized clone hN2Y showed slightly reduced affinity for the Beta variant compared to the original rabbit clone C-A11 (**Fig. 3** and **Table 2**), but affinity was increased to low pM binding K_D_ for the light chain engineered clone LxC1-G10. LxC1-G10 possessed 24 pM K_D_ affinity for BA.1, 5-fold improved over hN2Y. However, LxC1-G10 binding affinity was decreased nearly 100-fold to 2.1 nM for BQ.1.1 compared to 24 pM for BA.1.

**Fig. 3.**
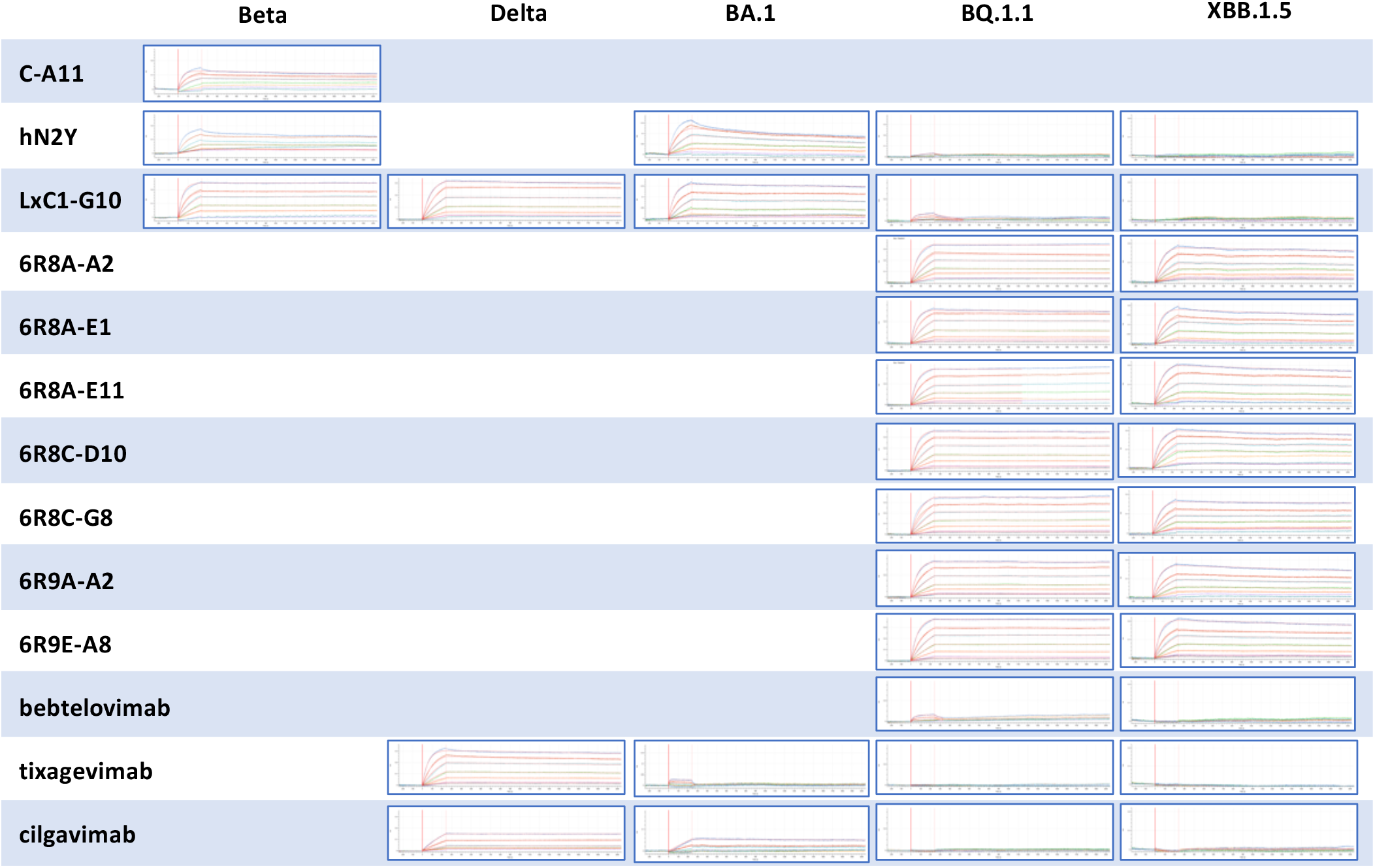
BLI affinity measurement shows strong binding to BQ.1.1 and XBB.1.5 for engineered IgGs. In each assay, biotinylated spike protein trimer was captured to the streptavidin-coated sensor surface and dipped into a 7-fold dilution series of IgG at a starting concentration of 10 nM. Binding was monitored for 4 minutes, followed by dissociation monitored for 30 minutes. For each variant tested, each experiment is shown with the same scale y-axis. Both the humanized lead candidate hN2Y and the light chain engineered clone LxC1-G10 lost binding activity for later SARS-CoV-2 variants, but all seven 6R8/6R9 clones showed strong binding to BQ.1.1 and XBB.1.5. Therapeutic mAbs tested under the same conditions showed weak or no binding to the latest Omicron variants.

**Table 2.**
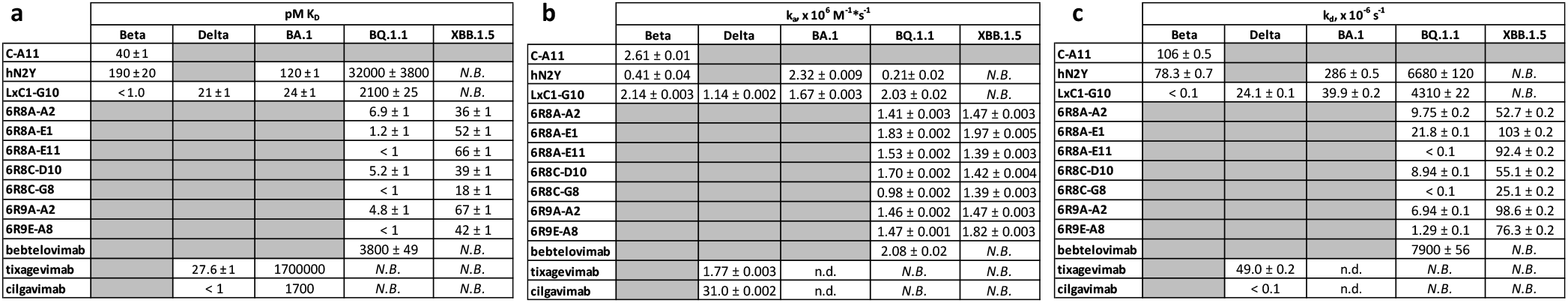
Affinity and kinetic values derived from BLI measurements. BLI traces were globally fit to a 1:1 binding model to derive **a**, K_D_ affinity values, **b**, on-rates, and **c**, off-rates. Some IgG affinities were below the detection limit of the assay (< 1 pM) due to exceptionally slow off-rates even when measured for 30 minutes. For some weakly binding clones, affinity and/or kinetic values could not be determined. *N.B*., non-binding; n.d., not derived.

The seven 6R8/6R9 IgGs possessed 7 pM or lower K_D_ for BQ.1.1 and below 70 pM K_D_ for XBB.1.5. Binding affinity for these mAbs was increased nearly 10,000-fold (for BQ.1.1) or above undetectable levels (for XBB.1.5) compared to parent clone hN2Y. The difference in binding affinity for BQ.1.1 is primarily due to a nearly 1000-fold reduction in off rate, with hN2Y possessing 10^-3^ s^-1^ off-rate while 6R8/6R9 clones showed 10^-6^ s^-1^ off rates.

In the assay, Bebtelovimab showed weak binding to BQ.1.1 and no binding at all to XBB.1.5. Evusheld is an antibody cocktail of Tixagevimab and Cilgavimab first given Emergency Use Authorization by the US FDA in December 2021, though the agency released a notice in January 2023 warning that Evusheld was not expected to neutralize XBB.1.5. Both Tixagevimab and Cilgavimab showed no binding to BQ.1.1 or to XBB.1.5 variants. In contrast, our engineered IgGs show low pM binding K_D_ for these latest SARS-CoV-2 Omicron variants currently circulating.

## Discussion

We have detailed the engineering and activity of broadly neutralizing mAbs that show potent efficacy against the latest Omicron circulating variants XBB.1.5 and BQ.1.1. Uniquely, instead of identifying a novel clone from infected or immunized samples, we rather iteratively engineered a single IgG, originally developed for neutralization of the Wuhan-Hu-1 strain, to broaden activity.

Our humanized clone hN2Y was derived from a rabbit lead candidate. Rabbits are an excellent source of potent antibodies due to the exceptionally high CDR sequence diversity of their immune repertoire by use of somatic gene conversion^25^. Despite this diversity, humanization of rabbit clones is relatively straightforward due to their restricted use of a limited number of germline genes^26^. C-A11 rabbit mAb possesses a 19 amino acid CDR H3. Long CDR H3 loops, though typically defined as ≥24 amino acids, are a common feature of broadly neutralizing antiviral mAbs^27^. Though not technically classified as ‘long’, this feature may have contributed to its potent neutralization of early variants through Delta. After humanization, hN2Y showed almost no change in affinity or neutralization activity compared to C-A11 and showed even stronger neutralization of some variants including BA.1 and BA.2.

As observed for many other therapeutic antibodies developed prior to December 2021, the first Omicron variant BA.1 reduced the potency of hN2Y. As detailed above, LxC1-G10 was selected by panning a combined light chain library on the Beta variant. The Beta variant only contains spike protein mutations K417N/E484K/N501Y. Nearly all Omicron subvariants retain these three mutations but additionally possess many others. LxC1-G10 showed sub-pM affinity for the Beta variant, >100-fold increased compared to hN2Y. This increased binding was sufficient to improve neutralization of early Omicron subvariants BA.1, BA.2, BA.2.75, BA.2.3.20, and BN.1, despite those variants possessing many additional mutations within the spike protein. LxC1-G10 shares an identical heavy chain as hN2Y, but possesses over 14 amino acid differences distributed across all light chain CDRs. Light chain CDR mutations were thus sufficient to confer significant improvement in binding affinity and neutralization potency.

However, LxC1-G10 lost activity against later Omicron subvariants from the BA.2 lineage that includes XBB.1.5 and from the BA.4/BA.5 lineage that includes BQ.1.1, likely primarily due to the presence of a mutation at F486. F486 mutations are found in all later Omicron variants and have been linked to immune evasion^1^. As detailed above, further engineering was performed by panning a combined light and heavy chain library on later Omicron variants. Seven strong candidates stood out from this selection, 6R8A-A2, 6R8A-E1, 6R8A-E11, 6R8C-D10, 6R8C-G8, 6R9A-A2, and 6R9E-A8. These show exceptional potency against all SARS-CoV-2 variants tested. Affinity was improved to single digit or sub-pM for BQ.1.1 and mid pM for XBB.1.5 for all seven mAbs. Compared to both hN2Y and LxC1-G10, 6R8/6R9 mAbs possess two mutations in CDR H3 (which is identical among all 6R8/6R9 clones) and 7-10 mutations in CDR H1. Compared to LxC1-G10, 6R8/6R9 mAbs possess 11-13 mutations across all light chain CDRs. Extensive mutagenesis across all CDRs except H2 was thus required to confer the increased neutralization activity and binding affinity observed.

Both our early lead candidate hN2Y (COVIC-359) and LxC1-G10 (COVIC-362) were submitted to the Coronavirus Immunotherapy Consortium (CoVIC) at La Jolla Institute of Immunology for analysis and testing against over 350 other therapeutic antibody candidates^28,29^. Both antibodies showed nearly 100% protection against cell infection in pseudovirus neutralization assay for Beta, Delta, BA.1, BA.1.1, and BA.2 variants, and showed efficacy with live Wuhan-Hu-1 virus in both neutralization assay and during *in vivo* mouse challenge. Both were binned into the same epitope class, RBD-2a, characterized by bivalent intra-spike binding, where both arms of the IgG bind to two spike units within a single trimer. This binding motif was further validated by cryo-EM, in which LxC1-G10 was observed to bind to both spike units in the ‘up’ conformation, a conformational state that exposes the ACE2 receptor binding site. The increased avidity conferred by bivalent binding likely contributes to the strong neutralization activity and high affinity for 6R8/6R9 mAbs.

We have characterized an antibody panel that outperforms all existing FDA-approved antibody therapeutics against the latest circulating SARS-CoV-2 strains. This study validates our unique STage-Enhanced Maturation (STEM) platform. Though our lead humanized clone showed no binding to or neutralization of XBB.1.5, we successfully recovered potency by engineering through repeated sampling and combination of CDRs progressively against SARS-CoV-2 variants. The ability to recover neutralization activity from a single lead mAb clone demonstrates the exceptional CDR sequence diversity that can be obtained by applying an iterative selection and library creation strategy. By continuing to engineer a single mAb, rather than repeating the discovery phase, lead time was significantly reduced. *In vitro* affinity maturation of a single neutralizing antibody has the potential to be far more efficient compared to discovery campaigns reliant on *in vivo* affinity maturation by successive vaccination and/or infection. In patient samples, broadly neutralizing antibodies must be identified from a large background of non-neutralizing anti-spike protein antibodies, while affinity maturation of a neutralizing antibody allows for fine tuning around a known neutralizing epitope. Additionally, filters can be included during phage panning, including heat treatment to remove unstable clones^30^ and negative subtraction on bacculovirus particles to remove polyreactive clones, to ensure that the final resulting candidates retain good developability profiles. This approach can be adapted for future therapeutic mAb development against SARS-CoV-2 or other viruses, even to allow for rapid development of broadly neutralizing therapeutic mAbs in real-time.

## Online Methods

### Preparation of recombinant proteins

Spike trimers were prepared in-house for use in all assays. SARS-CoV-2 variants were made by overlap PCR and cloned to pCAGGS vector for production. Spike trimers include polybasic cleavage site mutation/deletion and solubilizing mutations K986P/V987P as well as a trimerization motif and polyhistidine tag^31^. Following large-scale purification of DNA, 300 μg of DNA was transfected to HEK293T cells, with media harvested by centrifugation and filtration approximately 7 days post transfection. His-tagged spike trimer was purified using a Ni-NTA column: the spike trimer was washed with 50 mM imidazole then eluted with 100 mM and 250 mM imidazole, followed by overnight dialysis in DPBS. Quality and purity were assessed by SDS-PAGE gel and activity was assessed by testing for binding to ACE2-Fc by ELISA. In-house biotinylated trimers were prepared for BLI studies using EX-Link™ NHS-LC-Biotin (ThermoFisher A39257) followed by purification with 7K MWCO Zeba™ Spin Desalting Columns (ThermoFisher 89882). Spike trimers used in the study include Wuhan-Hu-1 (GenBank MN908947; Abwiz Bio Cat. #2720) as well as variants Beta (Abwiz Bio Cat. #2652), Delta (Abwiz Bio Cat. #2611), BA.1 (Abwiz Bio Cat. #2672), BA.2 (Abwiz Bio Cat. #2460), BA.2.75 (Abwiz Bio Cat. #2664), BA.2.75.2 (Abwiz Bio Cat. #2676), BA.2.3.20 (Abwiz Bio Cat. #2660), BN.1 (Abwiz Bio Cat. #2696), XBB (Abwiz Bio Cat. #2700), XBB.1 (Abwiz Bio Cat. #2648), XBB.1.5 (Abwiz Bio Cat. #2712), BA.4.6 (Abwiz Bio Cat. #2692), BA.5 (Abwiz Bio Cat. #2688), BQ.1 (Abwiz Bio Cat. #2704), BQ.1.1 (Abwiz Bio Cat. #2668), and BF.7/BA.5.2.6/BF.11 (Abwiz Bio Cat. #2708).

### Immunization of rabbits

Three New Zealand White rabbits were immunized with a recombinant protein including the receptor binding domain (RBD) of SARS-CoV-2 (Wuhan-Hu-1 strain). The immune sera were tested by ELISA using spike trimer, and the rabbit that showed the best titer was selected for library construction.

### Library construction

The bone marrow and spleen cells were collected and homogenized in TRI reagent (Molecular Research Center). Total RNA was isolated from the homogenate according to the manufacturer’s protocol. Messenger RNA was purified using mRNA purification kit (Macherey-Nagel) according to the manufacturer’s protocol. First strand cDNA was synthesized using PowerScribe MMLV Reverse Transcriptase (Monserate Biotechnology Group). First strand cDNA was engineered using a method fully described in the US patent 9,890,414. Briefly, the first strand cDNA was digested with a restriction endonuclease in the presence of an oligonucleotide and an adapter oligonucleotide with an artificial sequence on its 5’ end was ligated to the 5’ end of the digested cDNA using Taq DNA Ligase (New England Biolabs). Second strand cDNA was synthesized with framework region 1 specific primers with the same artificial sequence on their 5’ end. The finished 2nd strand cDNA was amplified by PCR with a single primer with the artificial sequence. The amplified products were purified by column and digested with restriction endonucleases. The digested fragments were run on an agarose gel and the appropriate sizes were purified. The light chain fragments were ligated with the phagemid vector and transformed into NEB 10-beta cells (New England Biolabs). The cells were grown overnight in the presence of carbenicillin and DNA was purified using Maxi DNA tips (Macherey-Nagel) according to the manufacturer’s protocol. The DNA was digested and run on a gel. The DNA was purified from the gel and ligated with the heavy chain fragment. NEB 10-beta cells were transformed with the ligation reaction and grown overnight in the presence of carbenicillin. DNA was purified using Maxi DNA tips (Macherey-Nagel) according to the manufacturer’s protocol.

### Phage panning and Fab screening

Library DNA was transformed into XL1-Blue cells (Agilent) and grown in the presence of carbenicillin and tetracycline. M13KO7 Helper Phage (New England Biolabs) was added and phage production was induced in the presence of 1 mM Isopropyl β-D-1-thiogalactopyranoside (IPTG) and kanamycin overnight. The culture was spun down and phage was precipitated with 4% PEG-8000/3% NaCl on ice. The precipitated phage was added to the wells coated with 100 μL of recombinant spike trimer at 37°C for 1 hour. The unbound page was washed with PBS and the bound phage was eluted twice with 0.1 M HCl (adjusted with glycine to pH2.2)/bovine serum albumin (BSA) 1 mg/mL) and neutralized with 2 M Tris. Freshly grown ER2738 cells (New England Biolabs) were infected with the eluted phage and grown in the presence of carbenicillin and tetracycline. M13KO7 Helper Phage (New England Biolabs) was added and phage production was induced in the presence of 1 mM IPTG and kanamycin overnight. The panning consisted of 4 rounds to enrich for specific binders. A portion of cells infected with the eluted page was used to make colonies on LB agar plates containing 100 μg/mL of carbenicillin for the screening of individual clones. The colonies were picked and added to 96-well culture plates with 1 mL of Super Broth medium and 50 μg/mL of carbenicillin. The culture plates were incubated at 37 °C and 1 mM IPTG was added to induce soluble Fab production at 30 °C. For Fab ELISA, microtiter wells were coated with 100 μL of spike trimer at 1-2 μg/mL in PBS and blocked with 200 μL of 1% BSA/PBS. The culture plates were spun down and the supernatant containing soluble Fab was added to the wells. The bound Fab was detected with peroxidase conjugated goat antirabbit IgG F(ab’)2 (Thermo Fisher Scientific). Positive clones were PCR amplified and sequenced.

### Surrogate virus neutralization tests (sVNT) of crude Fab

Surrogate virus neutralization tests (sVNTs) were performed according to the protocol described previously^22^ with some modifications. Microtiter wells were coated with 100 μL of ACE2-Fc (Abwiz Cat. #2566) at 2 μg/mL in PBS and blocked with 200 μL of 1% BSA/PBS. Fab supernatants were mixed 1:1 with a recombinant spike trimer at RT and added to the wells. The bound spike trimer was detected with peroxidase conjugated rabbit anti-his tag antibody (Jackson ImmunoResearch 300-035-240), and clones showing strong blocking were PCR amplified for sequencing of light and heavy chains.

### IgG production and humanization of rabbit antibody

Light and heavy chains of clone rabbit C-A11 were PCR amplified, digested, and cloned into a bi-cistronic rabbit IgG vector for expression. All antibodies and recombinant ACE2-Fc (Abwiz Bio Cat. #2566) were produced by transient transfection of HEK293T cells with media harvested by centrifugation and filtration approximately 7 days post transfection. IgGs were purified by protein A column (Cytiva) followed by overnight dialysis in DPBS. CDRs of C-A11 rabbit antibody were grafted into selected human germline genes (IGHV3-23*04 and IGKV-39*01) to create a humanized clone hN2Y. An expanded structural definition of CDRs was used, based on observed diversity from an internal database of rabbit antibodies. Following gene synthesis, hN2Y light and heavy chain genes were cloned into a bi-cistronic human IgG vector for production as above.

### Surrogate virus neutralization tests of purified IgG (sVNT)

Microtiter wells were coated with 100 μL of ACE2-Fc (Abwiz Cat. #2566) at 2 μg/mL in PBS and blocked with 200 μL of 1% BSA/PBS. Purified IgG were mixed 1:1 with a recombinant spike trimer at RT and added to the wells. The bound spike trimer was detected with peroxidase conjugated anti-his tag antibody (Jackson ImmunoResearch 300-035-240). All assays were performed in triplicate. Raw ELISA values were normalized to 6 replicate positive wells omitting IgG and to 3 negative wells omitting both IgG and spike trimer. Graphpad Prism was used to prepare graphs and to derive inhibition of spike trimer binding to ACE2 (IC_50_) values. Bebtelovimab, Tixagevimab, Cilgavimab, and S2X324 were purchased from Proteogenix (Schiltigheim, France). S728-1157 light and heavy chain genes were synthesized by Twist Bioscience (San Francisco, CA) and cloned into our IgG expression vector for expression and purification as described above.

### Construction of single CDR mutant libraries

Six single CDR mutant libraries were designed based on amino acid usage by structurally characterized human antibodies sharing similar canonical CDR structures among a subset that possessed identical germline genes. This design strategy further humanizes the amino acid usage in the CDRs and helps to avoid unwanted mutations that could negatively affect the structure and folding of the antibodies while constraining library size to a diversity that could be sampled using our phage libraries. Mutagenic oligos were synthesized and used to construct separate CDR libraries by fragment amplification and overlap PCR using hN2Y gene as a template. Following digestion, ligation was performed at large scale using 10 μg digested vector DNA. Library quality was assessed by titering to ensure low background of undigested vector DNA and by Sanger sequencing of 12 random colonies to ensure a sufficient proportion of full-length clones.

### Phage panning and Fab screening of single CDR mutant libraries

Single CDR mutant libraries were panned on Wuhan-Hu-1 spike trimer to remove non-productive clones and to enrich all possible CDR mutants that retain binding to the spike trimer. Phage panning was performed for four rounds as described above, using 5 μg/mL of trimer coated to ELISA wells and by increasing stringency following each round of selection. Fabs were screened similarly as described above and tested for binding to Wuhan-Hu-1 spike trimer. Binding clones from each library were PCR amplified and sequenced to determine the diversity of the selected clones.

### Library construction, panning, and screening of separate combined light and combined heavy chain libraries

Round 4 output phage from pre-selected light chain single CDR mutant library panning were used to amplify targeted CDR regions and combined by overlap PCR (Lx); this library was then ligated to a vector containing the wild-type hN2Y heavy chain. The phage from pre-selected heavy chain single CDR mutant libraries were similarly amplified and combined by overlap PCR (Hx); this library was then ligated to a vector containing the wild-type hN2Y light chain. Large scale library construction and QC was performed as detailed above. The combined light chain library was panned on the Beta spike trimer, and the combined heavy chain library was panned separately on Wuhan-Hu-1, Alpha, Beta, and Epsilon spike trimers, with increasing washing stringency each round of selection. Fabs were screened similarly as described above and tested for binding to the respective spike trimer used for selection. Binding clones from each library were PCR amplified and sequenced. LxC1-G10 clone was identified from the combined light chain library and was cloned to IgG format and characterized as described above.

### Library construction, panning, and screening of a single combined light and heavy chain library

Round 4 output phage from pre-selected combined light chain library panned on Beta spike trimer and from preselected combined heavy chain libraries panned on Wuha-Hu-1, Alpha, Beta, and Epsilon spike trimers were amplified by PCR and cloned into the phagemid vector as one library. Large scale library construction and QC was performed as detailed above. This library was panned for six rounds on BA.5 followed by two rounds on BA.2.75 and one round on either BQ.1.1 or XBB spike trimers. Fabs were screened similarly as described above and tested for binding to the respective spike trimer used for selection. Binding clones from each library were PCR amplified and sequenced. Clones possessing 6R8 or 6R9 prefixes were identified from this screening and were converted to IgG format for characterization as described above.

### Biolayer interferometry affinity measurement

Biotinylated spike trimers were prepared in-house as described above excepting BA.1 (ACROBiosystems SPN-C82Ee). BLI was performed using Octet Red96e with data collection at 30°C and sample shaking at 1000 rpm during all steps. All IgGs and spike trimers were prepared in running buffer containing 2% BSA, 0.002% Tween-20, PBS, pH 7.4. Following 4 minute baseline, 3 μg/mL of biotinylated trimer was captured to streptavidin sensors (Sartorius 18-5019) for 4 minutes followed by an additional 4 minute baseline. IgG binding was monitored for 4 minutes followed by 30 minute dissociation in fresh buffer only wells. Each IgG was tested in 1:2 titration series from 10 to 0.16 nM. A reference sensor omitting IgG was included in each assay and used for subtraction. All data were globally fit to 1:1 binding model to derive kinetic and affinity values, only omitting individual traces or times due to sensor error or low signal.

## Supplementary Data

**Fig. S1.**
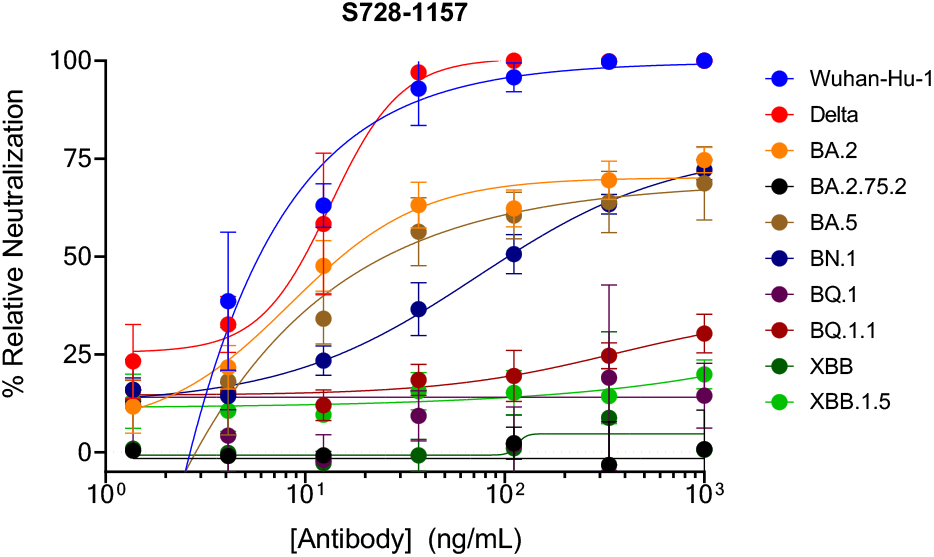
sVNT of S728-1157 mAb. S728-1157 showed potent neutralization of Wuhan-Hu-1 and Delta but reduced neutralization of BA.2, BA.5, and BN.1. Nearly no neutralization was observed for BA.2.75.2, BQ.1, BQ.1.1, XBB, or XBB.1.5.

**Fig. S2.**
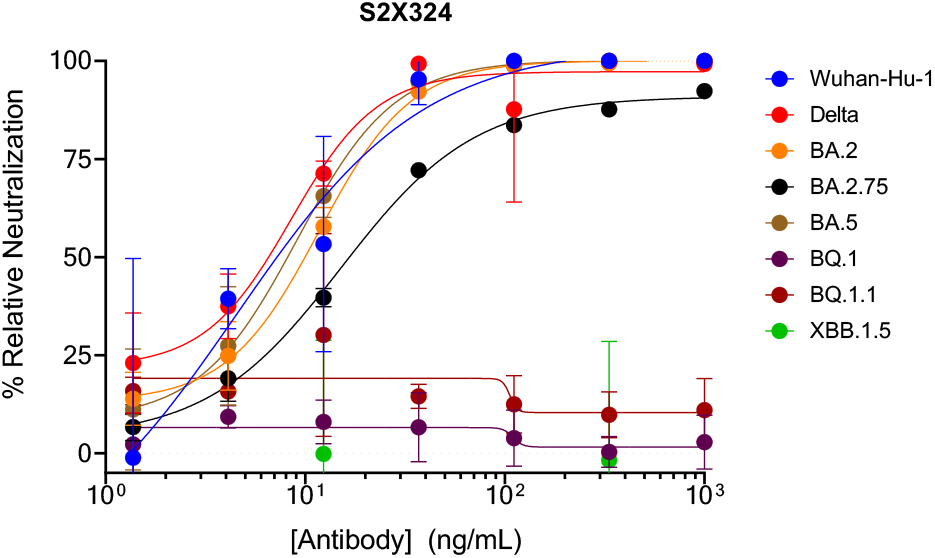
sVNT of S2X324. S2X324 showed potent neutralization of Wuhan-Hu-1, Delta, BA.2, BA.2.75, and BA.5. No neutralization was observed for BQ.1, BQ.1.1, or XBB.1.5.

